# Impact of the pear psyllid *Cacopsylla pyri* host instar on the behavior and fitness of the parasitoid *Trechnites insidious*

**DOI:** 10.1101/2020.11.09.374504

**Authors:** Guillaume Jean Le Goff, Jeremy Berthe, Kévin Tougeron, Benoit Dochy, Olivier Lebbe, François Renoz, Thierry Hance

**Author notes:** Corresponding author: Guillaume Jean Le Goff (E-mail address).

## Abstract

1. Pear is one of the most important fruit crops of temperate regions. The control of its mains pest, *Cacopsylla pyri*, is still largely based on the use of chemical pesticides, with all that this implies in terms of negative effects on the environment and health.
2. Within the context of integrated pest management, innovative and ecologically sustainable strategies must be developed for. Although *Trechnites insidiosus* is the most abundant parasitoid of *C. pyri*, its biology and its potential as a control agent have been little studied.
3. In this paper, we conducted experiments to evaluate the behavior of the specialist parasitoid *T. insidiosus* when exposed to different larval instars of the pear psyllid *C. pyri*, and to assess the quality of the next generation of parasitoids.
4. We found that although *T. insidiosus* accepts all host instars for oviposition, the third and fourth instars were the most suitable host in terms of behavioral acceptance and progeny development.
5. Our study is a first step for further studies on the interaction between psyllids and parasitoids and provides evidence that the specialist parasitoid *T. insidiosus* is a promising candidate for biological control strategies of the pear psyllid *C. pyri*.

## Introduction

Pear (*Pyrus communis* L.) is one of the major cash crops in temperate climates, with 1,381,923 hectares dedicated to its cultivation and approximately 24 million tons of pear produced in orchards in 2018 (“FAOSTAT,” 2020). The European pear psylla *Cacopsylla pyri* L. (Hemiptera: Psyllidae) is the most important pest of European pear trees. It infests all commercial pear tree varieties, is distributed throughout the pear cultivation area, and therefore presents a serious threat to pear production and causes heavy economic losses (Civolani, 2012; Dupont and Strohm, 2020). Both immature and adult psyllids feed on phloem sap and cause direct damages to pear trees and yield losses by subtracting nutrients and weakening the host plant. *C. pyri* also cause indirect damages by spreading fungi, such as sooty molds, mostly caused by a high honeydew excretion (Civolani, 2012; Sanchez and Ortin-Angulo, 2012). These pests also transmit various phytopathogens such as the phytoplasma Candidatus *Phytoplasma pyri* (Seemüller and Schneider, 2004), which is responsible of pear decline disease, reducing tree vigor (Civolani, 2012).

Control of *C. pyri* is currently mainly based on Integrated Pest Management (IPM), which relies on a mixture of natural and chemical control strategies (Civolani, 2012). However, excessive use of non-selective chemicals alone decreases the effectiveness of spray treatments over time due to the development of biotic resistance in psyllid populations (Buès et al., 2003; Erler, 2004; Civolani et al., 2007). With the banning of an increasing number of phytochemicals and the increasing public demand for organic food production, it is important to develop new approaches for the control of psyllids that are more environmentally and human health friendly. Biological control and the use of beneficial insects are already used as part of IPM strategies in pear orchards. For example, several bug species of the Anthocoridae family (Hemiptera) (e.g. *Anthocoris nemoralis*) are generalist predators of the pear psylla in orchards. The natural the presence of these generalist predators in pear orchards is not sufficient to regulate psyllid populations below sustainable economic threshold (Booth, 1992; Erler, 2004; Civolani, 2012), but mass releases of anthocorids into orchards have been shown to be effective in reducing pear psyllid populations (Sigsgaard et al., 2006a,b). However, the use of these generalist predators on a large scale faces many limitations, including the high cost of mass releases, the high pesticide sensitivity of anthocorids, and the lack of targeted control with a broad spectrum of insects that can be targeted by these predators (Booth, 1992; Civolani, 2012; Emami et al., 2014). To overcome the weaknesses of the currently used control methods, it is important to find more specific biological control strategies to meet the current demand for fruit produced in a more environmentally friendly and healthy way.

The use of specialist parasitoids are promising alternatives or complements to the use of generalist predators, due to their host-specificity, foraging capacity, high fecundity rate, and lack of negative side-effects in the environment. The parasitofauna of *C. pyri* is quite diversified and several species have been reported in pear orchards, including *Trechnites insidiosus, Prionomitus mitratus* (Dalman), *P. tiliaris* (Dalma,), *Endopsylla sp*., *Psyllaephagus procerus* Marcet*, Syrphophagus ariantes* (Walker)*, Syrphophagus taeniatus* (Fo□rster) and *Tamarixia* sp (Armand et al., 1990, 1991; Erler, 2004; Guerrieri and Noyes, 2009; Jerinic-Prodanovic et al., 2010). However, no attempts have been reported to rear these species for mass production, and little information is available on their biology, general ecology, and potential effects in a biological control context (Tougeron et al., 2021). Of these parasitoids, *T. insidiosus* is consistently cited as the most abundant species in the pear orchards (Armand et al., 1991, 1990; Avilla and Artigues, 1992; Booth, 1992; Bufaur et al., 2010; Erler, 2004; Herard, 1985; Miliczky and Horton, 2005; Sanchez and Ortin-Angulo, 2012), even though it exhibits significant susceptibility to chemical treatments (Burts, 1983; Lacey et al., 2005; Sanchez and Ortin-Angulo, 2012) and relatively high levels of hyperparasitism (McMullen, 1966; Armand et al., 1991, 1990; Sanchez and Ortin-Angulo, 2012). It is a koinobiont parasitoid that has interesting attributes for IPM of pear psyllids. First, it has a wide activity window, from early April to late November, which means it can be active at fairly low temperatures (Armand et al., 1991, 1990; Bufaur et al., 2010; Dupont and Strohm, 2020; Herard, 1985; Oudeh et al., 2013). *T. insidiosus* also has a first generation free of hyperparasitism (Armand et al., 1991, 1990). This parasitoid is therefore capable of significant and early activity at the first generation of pear psyllids, before the arrival of massive predators. *T. insidiosus* has been purposely introduced in California as a biological control agent to limit populations of introduced psyllid pests (Retan and Peterson, 1982; Unruh et al., 1995; Guerrieri and Noyes, 2009), but little information is available on its establishment success and effectiveness in controlling psyllids. The few field studies have revealed peak parasitism levels that vary between 30 to 100% depending on location (Bufaur et al., 2010; Erler, 2004; Jaworska et al., 1998; Oudeh et al., 2013), suggesting effective control of psyllid populations by *T. insidiosus* (Talitski, 1996 in Unruh et al., 1994).

To date, no experimental studies have been conducted to identify the parasitic behavior of *T. insidiosus* towards the pear psyllid *Cacopsilla pyri*. In this study, we conducted experiments to determine the set of behaviors that female *T. insidiosus* exhibit towards *C. pyri* as well as the physiological responses of *T. insidiosus* as a function of host developmental instar. The quality of the next generation parasitoids was assessed by measuring their developmental time, fecundity, size, and sex ratio, which are commonly used as proxies for assessing parasitoid fitness (Colinet et al., 2005). Previous studies have reported *T. insidiosus* females to preferentially lay eggs in the fourth and fifth larval instars of pear psyllids (Armand et al., 1991, 1990; Booth, 1992), while others have reported that they primarily oviposit in the first and second larval instars of psyllids (McMullen, 1966). Our study assumes that *T. insidiosus* females are capable of ovipositing in all larval development instars of *C. pyri*, with a preference for the more mature instars that would be more suitable hosts from a nutritional standpoint. We also hypothesized that parasitoids emerging from the older developmental instars exhibit better fitness indicators than individuals developing in younger instars of the psyllid.

## Material and methods

### Biological models

The insects used for the experiments were initially collected from populations sampled in 2013 for *Cacopsylla pyri* and in 2016 for *Trechnites insidiosus* in the experimental pear orchard of Proefcentrum voor Fruitteelt, Sint-Truiden-Belgium. Populations were maintained in the laboratory on pear trees in standardized sequential rearing conditions, that allowed to control the instar and the age of the insects, under the following conditions: 24°C, 60% relative humidity, and 18:8 h photoperiod (Light:Dark).

### The influence of host development on parasitoid behavior

Groups of twenty psyllid larvae of the same developmental instar (first-, second-, third-, fourth-, and fifth-instar larvae) were placed on artificial diet feeders and were let settling for two hours. Artificial feeder setups consisted of 500 μL of a nutritive solution (supplied by Viridaxis SA, Belgium) placed on top of a 5.5 cm diameter Petri dish and spread with a piece of parafilm 5 cm square and stretched over the dish. The use of an artificial diet in the experiment allows to standardize the environment and avoids a potential influence of the host plant on the behavior of the parasitoids. The differentiation of larval instars was based on morphological criteria: the first three larval instars are creamy-yellow colored, while the fourth and the fifth instars vary between greenish-brown to dark-brown (Chang, 1977). The first instar larva is the same size as psyllid eggs, the second instar larva is twice as large, and the third instar larva has wing pads, which gradually grow in the fourth and fifth larval instars (Chang, 1977). A mated *T. insidiosus* female aged of less than 48 hours was then placed at the center of the set up and its behavior was recorded for thirty minutes with a Sony handycam (HDR XR200VE), during the afternoon. Ten replicates par larval instar were performed, using a naïve parasitoid female for each replicate. Using the event recorder software ODRec 3.0 (© Samuel Péan), we recorded and quantified: 1) the number of host-feeding behavior (when the parasitoid actively feeds on the host), 2) the time spent feeding on the host (expressed as a percentage of the total experimental time), 3) the time spent grooming (expressed as a percentage of the total experimental time), 4) the time spent walking (expressed as a percentage of the total experimental time), 5) the time spent resting (expressed as a percentage of the total experimental time), 6) the number of antennal contacts with the psyllid, and 8) the number of ovipositor insertions into the host (Albitar et al., 2016; Augustin et al., 2020). The host acceptance rate was calculated as the number of ovipositor insertions divided by the number of antennal contacts.

### The influence of host development on parasitism and offspring quality

After the behavioral bioassays, all psyllid larvae from a same replicate were placed on a same pear tree for fourteen days to await the formation of mummies (i.e., dead psyllids containing a developing parasitoid). We used in-vitro-cultivated pear trees (Williams cultivar) that were between one and two years old, and between 0.75 and 1 meter in height. Plants were obtained from Battistini Vivai (https://www.battisti-rebschule.it) and stored in individual cages in a climatic chamber at 24°C. After fourteen days, pear trees were checked daily for the presence of mummies and adult psyllids. Each mummy was then isolated in a falcon tube with a drop of honeydew until the parasitoid emerged. Development time was measured as the number of days from oviposition to adult emergence. Host suitability (number of mummies divided by the number of ovipositor insertions) was calculated for each host instar to determine the developmental instar that provides the best egg-to-adult development. The emergence rate was also calculated as the number of emerging adult parasitoid divided by the total number of mummies, for each treatment. Finally, the sex-ratio was calculated as the number of males divided by the total number of emerging individuals. Three days after emergence, parasitoids (males and females) were stored in a freezer at −20°C for future measurements of tibia size and egg load.

The size of the tibia was used as a proxy for individual body size. The left hind tibia of each emerging individual was measured using the software ImageJ 1.440 (Rasband, W.S., US National Institutes of Health, Bethesda, MD, USA). To estimate their egg load, each emerging female was squeezed from the abdomen beneath a cover slip on a microscope slide (Manfield and milles, 2002): the female was placed on an object blade with a small amount of water and crushed with a coverslip. To better extract the eggs, the pressure exerted on the coverslip started from the head towards the abdomen. Only ellipsoidal mature eggs (Figure 1) were counted.

**Figure 1:**
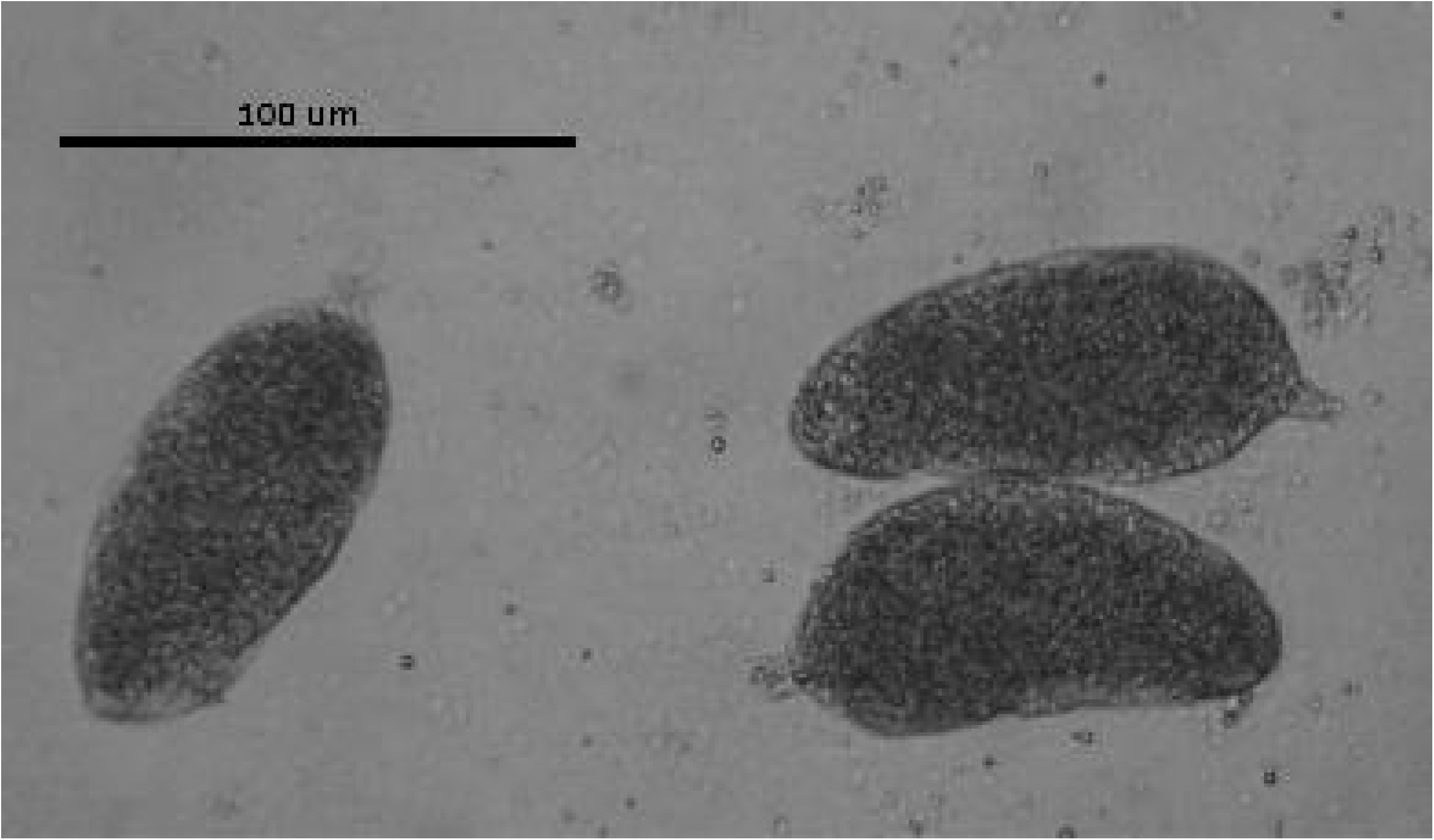
mature eggs of *Trechnites insidiosus*.

### Statistical analysis

Generalized linear models (GLMs) were fitted to the data to test the potential influence of host instar (explanatory variable, with five levels) on female parasitoids behaviors and emerging parasitoid quality. Response variables were the number of host feeding events (Poisson distribution), the time spent host feeding (Gaussian distribution), the time spent grooming (Gaussian distribution), the time spent walking (Gaussian distribution), the time spent resting (Gaussian distribution), the number of antennal contacts (Poisson distribution), the number of ovipositor insertions (Poisson distribution), the host acceptance rate (Gaussian distribution), the number of mummies (Poisson distribution), the host suitability (Gaussian distribution), the emergence rate (Binomial distribution), and the egg load (Poisson distribution).

We also used a GLM (Gaussian distribution) to test the potential influence of the sex, of the host instar, and of their interaction, on the tibia size and the development of emerging parasitoids. All significant GLMs were followed by Tukey post hoc tests to compare each level of the same factor (host instar and sex). In addition, Spearman correlation tests were performed between the tibia size and the egg load at the emergence of each female, for each host instar. Finally, using χ^2^ tests, we compared the experimental results of sex ratio obtained for each larval instar to a 50/50 theoretical sex ratio.

Statistical analyses were performed using R version 3.3.3 Copyright (C) 2016 The R Foundation for Statistical Computing for Mac. All tests were applied under two-tailed hypotheses, and the significance level was set at 0.05.

## Results

### The influence of host development on parasitoid behavior

The number of antennal contacts varied significantly with host developmental instar (χ^2^=800.30, DF=4, P<0.001). The minimum value was observed for *T. insidious* females exposed to second instar psyllids, while the maximum was observed for individuals exposed to third and fourth larval instars (Table 1) (S1). Regarding the average number of ovipositor insertions, it was significantly different between larval instars (χ^2^=443.92, DF=4, P<0.01): the second larval instar received a significantly lower number of ovipositor insertions than the third, fourth and fifth larval instars (Table 1) (S1). Host acceptance by parasitoids was significantly different between the different developmental instars (F=5.01, D=4, P<0.01), with acceptance for fifth instars lower than for all other developmental instars (Table 1) (S1).

**Table 1:** Mean total number and mean total duration ± standard deviation of the different observed behaviours, and number of replicates for each psyllid larval host stage. Different letters indicate significant differences to Tukey HSD tests.

|  | **Instar 1** | **Instar 2** | **Instar 3** | **Instar 4** | **Instar 5** |
| --- | --- | --- | --- | --- | --- |
| **Number of antennal contacts** | 17.50 ± 21.06 a<br>(n=10) | 9.60 ± 13.66 b<br>(n=10) | 32.10 ± 30.19 c<br>(n=10) | 31.70 ± 16.26 c<br>(n=10) | 16.00 ± 10.19 a<br>(n=10) |
| **Number of ovipositor insertions** | 10.00 ± 13.33 a<br>(n=10) | 4.70 ± 7.90 b<br>(n=10) | 14.80 ± 11.72 c<br>(n=10) | 17.80 ± 11.31 c<br>(n=10) | 4.00 ± 3.62 b<br>(n=10) |
| **Host acceptance** | 0.56 ± 0.27 a<br>(n=6) | 0.54 ± 0.31 a<br>(n=5) | 0.50 ± 0.17 a<br>(n=9) | 0.55 ± 0.15 a<br>(n=10) | 0.22 ± 0.11 b<br>(n=10) |
| **Time walking (%)** | 28.27 ± 21.72 ab<br>(n=10) | 18.49 ± 16.34 a<br>(n=10) | 36.06 ± 18.77 ab<br>(n=10) | 33.32 ± 6.91 ab<br>(n=10) | 42.26 ± 10.73 b<br>(n=10) |
| **Time resting (%)** | 16.73 ± 25.22 ab<br>(n=10) | 32.14 ± 31.67 a<br>(n=10) | 3.76 ± 6.14 b<br>(n=10) | 0.86 ± 1.53 b<br>(n=10) | 0.71 ± 1.84 b<br>(n=10) |
| **Number of host feeding events** | 0.00 ± 0.00 a<br>(n=10) | 0.10 ± 0.32 a<br>(n=10) | 0.10 ± 0.32 a<br>(n=10) | 0.10 ± 0.32 a<br>(n=10) | 0.00 ± 0.00 a<br>(n=10) |
| **Host feeding duration (%)** | 0.00 ± 0.00 a<br>(n=10) | 0.20 ± 0.62 a<br>(n=10) | 0.01 ± 0.04 a<br>(n=10) | 0.22 ± 0.70 a<br>(n=10) | 0.00 ± 0.00 a<br>(n=10) |
| **Grooming duration (%)** | 36.16 ± 19.97 a<br>(n=10) | 40.19 ± 16.67 a<br>(n=10) | 40.15 ± 21.65 a<br>(n=10) | 41.81 ± 11.04 a<br>(n=10) | 49.74 ± 11.01 a<br>(n=10) |

Host instar significantly influenced the time spent by parasitoids walking (F=3.19, DF=4, P<0.05). In the presence of fifth instar larvae, the parasitoid spent significantly more time walking, than in the presence of second instar larvae (Table 1) (S1). Host developmental instar had a significant impact on the time spent resting (F=5.50, DF=4, P<0.01). Parasitoids exposed to third, fourth and fifth instar larvae spent less time resting than those exposed to second instar larvae (Table 1) (S1).

The average number of host-feeding behaviors was very low; for each developmental instar about 1 out of 200 larvae were killed and then eaten by a parasitoid for an average duration of 0.10% of the total experimental time. No significant differences between developmental instars were observed for the occurrence (X^2^=0.19, DF=4, P=0.10) or the duration of host feeding (F=0.72, DF=4, P=0.58) (Table 1) (S1). Grooming accounted for an important part of the behavior expressed by the parasitoid over thirty minutes, and was expressed in a similar proportion in all developmental instars tested (around 42%) (F=0.90, DF=4, P=0.47) (Table 1) (S1).

### The influence of host development on parasitism and offspring quality

The average number of mummies was significantly different between developmental instars (X^2^=111.22, DF=4, P<0.001), with a higher mean number of mummies produced when parasitizing third- and fourth instar psyllids than first- and second instar psyllids (Table 2) (S1). Host suitability was significantly influenced by psyllid developmental instar (F=8.50, DF=4, P<0.001), with a lower suitability of first instar hosts than third instars hosts. Fifth instars showed a null suitability as they produced no mummies (Table 2) (S1). The emergence rate did not vary significantly with host developmental instar (χ^2^=44.463, DF=3, P>0.05). Of the 162 mummies obtained, 155 resulted in emergence. All mummies resulted in emergence when development occurred in the third instar (72/72), whereas when development of parasitoids occurred in first and fourth instar larvea, two mummies did not emerge (2/14 and 2/60, respectively) and for development in second instar larvae, three mummies did not emerge (3/16).

**Table 2:** Mean ± standard deviation of the different measured parameters of emerging parasitoids quality, and number of replicates for each psyllid larval host stage. Different letters indicate significant differences according to Tukey HSD tests.

|  | Instar 1 | Instar 2 | Instar 3 | Instar 4 | Instar 5 |
| --- | --- | --- | --- | --- | --- |
| **Number of mummies** | 1.33 ± 1.73 a<br>(n=10) | 1.44 ± 2.65<br>a (n=10) | 6.55 ± 4.61<br>b (n=10) | 5.78 ± 3.67 b<br>(n=10) | 0.00 ± 0.00 c<br>(n=10) |
| **Host suitability** | 0.15 ± 0.18 ac<br>(n=6) | 0.52 ± 0.28<br>ab (n=5) | 0.60 ± 0.36<br>b (n=10) | 0.40 ± 0.24<br>ab (n=10) | 0.00 ± 0.00 c<br>(n=9) |
| **Sex ratio** | 0.33 | 0.30 | 0.49 | 0.21 |  |
| **Female tibia size (mm)** | 0.36 ± 0.02 a<br>(n=9) | 0.33 ± 0.03<br>b (n=9) | 0.36 ± 0.02<br>a (n=35) | 0.36 ± 0.02 a<br>(n=46) | |
| **Male tibia size (mm)** | 0.33 ± 0.02 a<br>(n=3) | 0.32 ± 0.02<br>b (n=4) | 0.33 ± 0.03<br>b (n=36) | 0.34 ± 0.02 b<br>(n=12) | |
| **Egg load** | 19.77 ± 10.50<br>a (n=9) | 8.48 ± 6.75<br>a (n=9) | 11.44 ± 7.45 a<br>(n=35) | 11.30 ± 10.70<br>a (n=46) | |
| **Developmental time of females (days)** | 30.33 ± 2.65<br>a (n=9) | 22.33 ± 3.74<br>b (n=9) | 22.66 ± 2.83 bc<br>(n=35) | 21.89 ± 2.08<br>c (n=46) | |
| **Developmental time of males (days)** | 31.33 ± 1.53 a<br>(n=3) | 26.25 ± 7.23<br>b (n=4) | 22.86 ± 2.83 bc<br>(n=36) | 20.50 ± 1.93<br>c (n=12) | |

Parasitoids emerging from the first, second, and third instar larvae had a balanced sex ratio (χ^2^=0.5, P>0.5, χ^2^=0.8, P>0.4, χ^2^=1.48, P>0.2, respectively), whereas individuals emerging from the fourth instar larvae had a sex ratio that was largely skewed in favor of females (37 females for 8 males) (χ^2^=8.52, P<0.01) (Table 2) (S1). Tibia lengths of parasitoids differed significantly between the two sexes, males being smaller than females (0.33 mm vs. 0.35 mm, respectively), regardless of the host developmental instar (F=43.35, DF=1, P<0.001) (Table 2) (S1). Tibia length also varied with host developmental instar (F=3.33, DF=3, P<0.05). Male and female parasitoids from second instar larvae were in average smaller than those developed from other developmental instars (Table 2) (S1). No interaction was detected between sex and developmental instar factors (F=0.96, DF=3, P=0.41).

No impact of host instar was observed on female egg load, which showed an average of 10.94 ± 9.00 mature eggs over all experimental conditions (X^2^=549.71, DF=3, P>0.2) (Table 2) (S1). However, a significant correlation between the tibia length and female egg load was observed for females that developed in fourth instar hosts (Spearman’s R = 0.50, P <0.001, n=46), but not for females that emerged from the other developmental instars (R = 0.38, P> 0.05, n=9, R = 0.66, P> 0.05, n=9, and R = 0.12, P> 0.05, n=35, for instar 1, 2, and 3, respectively) (Figure 2). The developmental time of parasitoids was significantly different among host instars (F = 36.11, DF = 3, P<0.001). Parasitoid eggs laid in first instar hosts took longer to emerge from the mummies than parasitoid eggs laid in other developmental instars (Table 2) (S1). There was no significant difference in development time between sexes (F=0.14, DF=1, P=0.71), and no interaction occurred between the two factors (F=1.52, DF=3, P=0.21) (Table 2) (S1).

**Figure 2:**
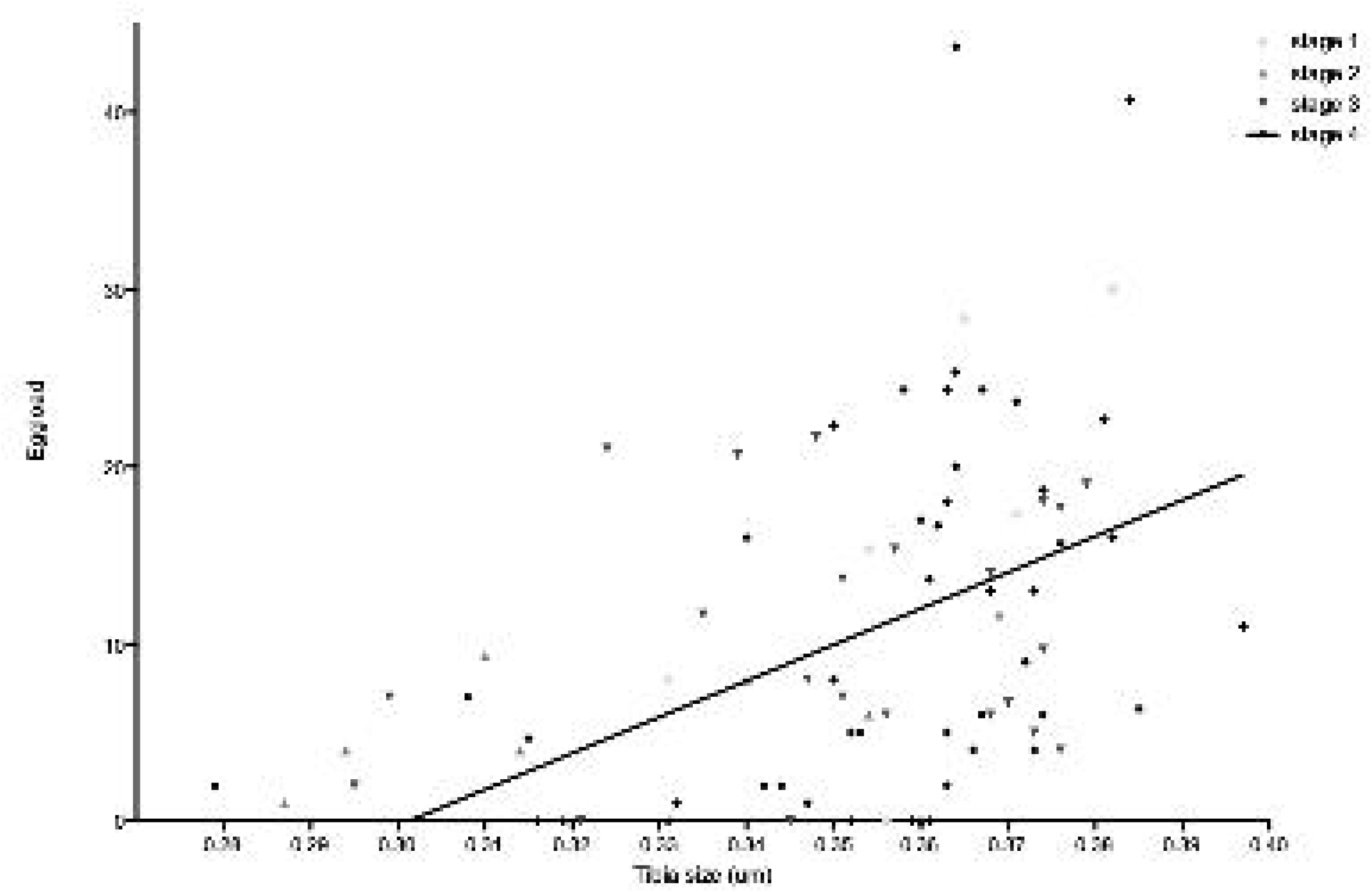
Correlation between the number of eggs per female from each developmental stage with tibia size.

## Discussion

According to Armand et al. (1991, 1990), and Booth (1992), *T. insidiosus* tends to oviposit in the fourth and fifth larval instars of the pear psyllid *C. pyri*, whereas McMullen (1966) suggest that this parasitoid oviposits mainly in the first three larval instars. In our study, we observed that this parasitoid accepted all host instars for oviposition. However, they had a lower acceptance rate for fifth instar hosts, in which no parasitoid could develop, and higher mummy production was found when eggs were laid in third and fourth instar larvae. Parasitoids spent more time resting and less time exploring the patch when exposed to the first two larval instars, and first instar hosts presented the lowest suitability for *T. insidiosus* of the instars in which mummy development was possible. Second instar hosts received few antennal contacts and as few ovipositor insertions as fifth instar hosts. First and second instar larvae together accounted for only 20 % of the total number of mummies produced in this experiment. In general, low parasitism rates of young host instars are associated with higher mortality of the host larvae, which are more susceptible to oviposition injuries from stinging and/or venom (Colinet et al., 2005). In addition, the mortality rate of young parasitized instars could be high because hosts are more likely to die between successive molts. All parameters taken together, we suggest that third and fourth instar psyllid larvae are the most suitable hosts for the development of *T. insidiosus*, both qualitatively and quantitatively.

We found that *Trechnites insidiosus* was more motivated to forage for hosts in presence of third, fourth and fifth instar larvae, with more time spent moving and less time spent resting than when exposed to the first and second instar larvae. Clues left by older psyllid larvae (e.g. honeydew, exuviae and chemical volatiles) could stimulate the locomotor activity of the parasitoid and thus increase its probability of finding hosts. This phenomenon was previously observed in the encytrid parasitoid *Psyllaephagus pistaciae* whose searching time, locomotion, antennal drumming, and ovipositor probing behaviors were increased by the presence of pistachio psylla honeydew (Mehrnejad and Copland, 2006). The antochorid predator *Orius sauteri* tends to forage more and to lay more eggs in the presence of the pear psylla *Cacopsylla chinensis* honeydew (Ge et al., 2019). Our results suggest that the amount and/or the quality of the clues present in the environment may be important stimulating cues for the parasitoid. Determining the nature of these clues influencing the exploratory behavior of *T. insidiosus* could be an additional step to unravel the factors driving the interactions between psyllids and parasitoids.

We found that third and fourth instar psyllids represent the most suitable hosts for oviposition, as 80 % of the mummies obtained in this experiment resulted from these two developmental instars. Although they are larger and therefore more difficult to handle by the parasitoid than first and second instar hosts, they appear to be the most suitable candidates for the female parasitoid, given the trade-off between the nutritional quality of the host and its behavioral and immune defense capabilities. From a biological control perspective, by parasitizing the fourth instars of *C. pyri*, *T. insidiosus* will mainly parasitized psyllid individuals that have escape to all other mortality factors at the end of their developmental cycle and just before the reproduction of the pest occurs. Such a feature gives the parasitoid a potentially important control efficiency on the population dynamics of its host with an immediate impact on the resulting imaginal population, and thus on the next generation of psyllids (Armand et al., 1991).

Our results showed a lower attraction and acceptance of *T. insidiosus* to fifth instar psyllids, as few antennal contacts and ovipositors insertion were performed, and no mummies were produced. Fifth instar larvae are probably too large and too advanced in ontogeny to allow proper development of *T. insidiosus*. Indeed, advanced larval instars of psyllids are able to escape the parasitoid more easily than the earlier developmental instars (Villagra et al., 2002). Such differences in escape behavior between host instars are commonly reported in aphid-parasitoid interactions, in which more mature hosts also generally have a higher capacity to encapsulate parasitoid eggs (Colinet et al., 2005). For example, it has also been shown that the last instar of the aphid *Toxoptera citricida* exhibits a greater immune response to parasitism than younger instars (Walker and Hoy, 2003). The absence of mummies for the fifth instar of psyllids could therefore be explained by a combination of behavioral and immune responses of the psyllid to parasitoid attack (Colinet et al., 2005). Fifth instar larvae of *C. pyri* are therefore completely unsuitable for parasitoid development.

Grooming accounts for nearly half of the activity of *T. insidiosus*, regardless of the host instar encountered. Psyllids, especially larvae, produce large amounts of honeydew (Civolani, 2012), which is highly concentrated in sugar (Le Goff et al., 2019). After an ovipositor insertion, residues of honeydew left on the cuticule probably promote bacterial and/or fungal infections on the parasitoid’s body. Selection likely favored individuals that spent a lot of time cleaning themselves (legs, ovipositor, antennae), because this behavior may not only help parasitoids to live longer, but also contributes maintain high levels of locomotor activity and host detection ability (Zhukovskaya et al., 2013). For psyllids, high honeydew production could also be a protection against parasitoids. Indeed, it has been observed that the honeydew of the pear psylla *Cacopsylla chinensis* limits the foraging behavior of its predators and could provide a physical defense for the psyllid (Ge et al., 2019). Moreover, *T. insidiosus* has been observed attempting to oviposit in in honeydew droplets, allowing time for psyllid larvae to escape. When attacking aphids, parasitoids also waste time manipulating and inserting their ovipositors into aphid exuviae (Muratori et al., 2008). Finally, the time *T. insidiosus* spends grooming itself is time that is not spent searching for a host. An experiment to analyze the behavior of the parasitoid when faced with exuviae of different instars and/or honeydew could be conducted to clarify the role that psyllid waste might play in its defense against parasitoids, in terms of the parasitoid’s time budget.

Regarding the parasitoid fitness indicators obtained in our experiments, parasitoids distribute their offspring in a balanced sex ratio in the first three host instars, while fourth instar psyllids were chosen by the parasitoid to lay a majority of females. It has already been shown that host size/instar can influence the sex ratio of parasitoid offspring: eggs leading to the emergence of females tend to be laid in large hosts (Bernal et al., 1997; Jervis and Kidd, 1986; Van Den Assem et al., 1982). This strategy is consistent with the host size distribution model, which assumes that the amount of resources available for parasitoid development determines its fitness (Charnov, 1976; Charnov and Skinner, 1985). Thus, it is more profitable for a female parasitoid to lay female eggs in large hosts that provide more resources (Jervis and Kidd, 1986), so they ultimately have a higher egg load. Our experiments were conducted with solitary females, but it would be interesting to test whether this species produces more males under conditions of competition for large hosts, as predicted by the theory of local mate competition (Hamilton, 1967).

The third and fourth larval instars of psyllids produced larger individuals than second instars, probably because these developmental instars exhibit more abundant reserves that allow for better growth of the parasitoid. More surprisingly, females that were laid in first instar hosts appeared to be as large as those that developed in third and fourth instar hosts. One mechanism that would explain these observations is that when an egg is laid in a first instar host, the egg does not begin to develop until the psyllid larva reaches a more advanced development instar (Colinet et al., 2005). This hypothesis is supported by the fact that individuals from a first instar host take longer to develop than individuals from other instars. It is also possible that parasitoid larvae develop less rapidly in such a host in order to keep it alive longer, thus ensuring completion of their development. These hypotheses could be confirmed by dissecting second, third and fourth instar larvae that were parasitized during the first development instar, and identifying the developmental instar of the parasitoid.

We observed a fairly low egg load in *T. insidiosus* females regardless of the host instar in which they developed, suggesting that this species is synovogenic and produces eggs throughout its life (approximately 20 days when fed under laboratory conditions (Berthe, 2018)). Furthermore, it is generally observed in parasitoids that the larger the female, the greater the egg load (Jia and Liu, 2018). In our study, this relationship was only observed for individuals from four instar hosts, confirming that this larval instar is the most suitable for parasitoid development. Yet, some large females did have no or few mature eggs. Yet, some large females had no or few mature eggs. It is possible that a stimulus such as psyllids honeydew, host feeding, or simply the presence of the host, is necessary to stimulate egg production (Aung et al., 2012; Pan et al., 2017).

Finally, although host feeding behavior was not affected by the host developmental instar, they may play an important role in the ecology of *T. insidiosus* and its interaction with the host. Host feeding is the consumption of host fluids exuding from oviposition wounds by the adult female parasitoid (Heimpel and Collier, 1996). This behavior has already been described in other encyrtidae species (Aung et al., 2012) but never in *T. insidiosus.* The number of host-feeding events that were observed in our experiments was very low, likely because the females we used were fed, hydrated and full of eggs. Therefore, their only optimal foraging strategy under these conditions was probably to lay as many eggs as possible. To better understand under what conditions host-feeding behavior is expressed, further experiments should be conducted on fertilized females with poor access to food, and/or with low egg loads. Such an experiment would highlight the trade-off between feeding to replenish their energetic reserves/egg load, and to lay eggs, when a femane meet a psyllid larvae. It is also possible that *T. insidiosus* is able to discriminate between a parasitized and a non-parasitized host, as is the case with most parasitoid species (van Alphen and Visser, 1990). Thus, a female arriving in a patch already visited by a conspecific would be more prone to express host-feeding behavior on a host parasitized by a competitor, and thus decrease the competition pressure encountered by its offspring. However, this hypothesis has yet to be tested.

The purpose of this study was to determine key elements in the interaction between pear psyllid *C. pyri* and the specialist parasitoid *T. insidiosus*, including the most profitable host instars for its development. We showed that third and fourth instar larvae are the most suitable hosts, both behaviorally and physiologically, for the parasitoid to produce high quantity and quality offspring. These results provide a basis for further studies on psyllid-parasitoid interactions, and are a first step to fill the data gap on the insect *T. insidiosus* in the literature, especially regarding its general biology and behavior, and open perspectives on the use of this parasitoid species in integrated pest management strategies against pear psyllids.

## Acknowledgments

This research was supported by the grant 12/1/7798 of the PPP call of the Walloon Region, by an additional grant from Viridaxis S.A. and by the Interreg-Proverbio project funded by SPW-Feder-Interreg (**https://www.interreg-fwvl.eu**). KT and FR were supported by the F.R.S.-FNRS. This publication is number BRC361 of the Biodiversity Research Centre.

## Author contribution

GJLG, JB, TH designed the study, JB made the experiments, GJLG and JB analyzed the data. BD, OL, GJLG caught the insect to start the rearing, maintained the rearing and the plant cultures. GJLG, KT, FR and TH wrote the manuscript. All authors contributed to manuscript improvement and gave their final approval for publication.

## Data availability statement

The datasets analyzed for the current study are available at the following DOI: https://doi.org/10.6084/m9.figshare.13187642.v1

## References

Albittar, L., Ismail, M., Bragard, C. & Hance, T. (2016) Host plants and aphid hosts influence the selection behaviour of three aphid parasitoids (Hymenoptera: Braconidae: Aphidiinae). European Journal of Entomology, 8.

Armand, E., Lyoussouf, A. & Rieux, R. (1991) Evolution du complexe parasitaire des psylles du poirier *Psylla pyri* et *Psylla pyrisuga* (Homoptera◻: Psyllidae). Entomophaga, 36, 287–294.

Armand, E., Lyoussoufi, A., Arcier, F.F. d’ & Rieux, R. (1990) Inter-relations entre les populations du psylle du poirier *Psylla pyri* (L.) (Hom., Psyllidae) et le complexe de ses parasitoïdes dans un verger traité du sud-est de la France. Journal of Applied Entomology, 110, 242–252.

Augustin, J., Boivin, G., Brodeur, J. & Bourgeois, G. (2020) Effect of temperature on the walking behaviour of an egg parasitoid: disentangling kinetic response from integrated response. Ecological Entomology, 45, 741–750.

Aung, K.S.D., Takasu, K., Ueno, T. & Takagi, M. (2012) Effect of Host–feeding on Reproduction in *Ooencyrtus nezarae* (Ishii) (Hymenoptera: Encyrtidae), an Egg Parasitoid of the Bean Bug *Riptortus clavatus*. Journal of the Faculty of Agriculture, Kyushu University, 57, 115–120.

Avilla, J. & Artigues, M. (1992) Parasitoides de Cacopsylla pyri (L.) (= *Psylla pyri* (L.)) presentes en una plantación comercial de peral en Lleida no sometida a tratamientos insecticidas. Bol. San. Veg. Plagas, 18, 133–138.

Bernal, J.S., Waggoner, M. & Gonzalez, D. (1997) Reproduction of *Aphelinus albidopus* (Hymenoptera: Aphelinidae) on russian wheat aphid (Hemiptera: Aphididae) hosts. European Journal of Entomology, 94, 83–96.

Berthe, J. (2018) Evaluation de la capacité de parasitisme de *Trechnites insidiosus* pour le contrôle biologique du psylle du poirier *Cacopsylla pyri* (Master Thesis).

Booth, S.R. (1992) The potential of endemic natural enemies to suppress pear psylla, *Cacopsylla pyricola* Förster, in the Hood River Valley, Oregon.

Buès, R., Boudinhon, L. & Toubon, J.-F. (2003) Resistance of pear psylla (*Cacopsylla pyri* L.; Hom., Psyllidae) to deltamethrin and synergism with piperonyl butoxide. Journal of Applied Entomology, 127, 305–312.

Bufaur, M., Harizanova, V. & Stoeva, A. (2010) Parasitoids of the pear sucker *Cacopsylla pyri* L. (Psyllidae) in Bulgaria - Morphology and biology. Presented at the Jubilee Scientific Conference with International Participation Traditions and Challenges of Agricultiral Education, Science and business, Agricultural Universitty - Plovdiv.

Burts, E.C. (1983) Effectiveness of a Soft-Pesticide Program on Pear Pests. Journal of Economic Entomology, 76, 936–941.

Chang, J. (1977) Studies on the susceptibility of pear trees to pear psylla, *Psylla pyricolla* Foerster (Homoptera: psylllidae).

Charnov, E.L. (1976) Optimal foraging, the marginal value theorem. Theoretical population biology, 9, 129–136.

Charnov, E.L. & Skinner, S.W. (1985) Complementary Approaches to the Understanding of Parasitoid Oviposition Decisions. Environmental Entomology, 14, 383–391.

Civolani, S. (2012) The Past and Present of Pear Protection Against the Pear Psylla, *Cacopsylla pyri* L. In *Insecticides* - *Pest Engineering* (ed. by Perveen, F.). InTech, pp. 385–408.

Civolani, S., Peretto, R., Caroli, L., Pasqualini, E., Chicca, M. & Leis, M. (2007) Preliminary Resistance Screening on Abamectin in Pear Psylla (Hemiptera: Psyllidae) in Northern Italy. Journal of Economic Entomology, 100, 1637–1641.

Colinet, H., Salin, C., Boivin, G. & Hance, Th. (2005) Host age and fitness-related traits in a koinobiont aphid parasitoid. Ecological Entomology, 30, 473–479.

Dupont, S.T. & Strohm, C.J. (2020) Integrated pest management programmes increase natural enemies of pear psylla in Central Washington pear orchards. Journal of Applied Entomology, 144, 109–122.

Emami, M.S., Shishehbor, P. & Karimzadeh, J. (2014) The influences of plant resistance on predation rate of *Anthocoris nemoralis* (Fabricius) on *Cacopsylla pyricola* (Förster). Archives of Phytopathology and Plant Protection, 47, 2043–2050.

Erler, F. (2004) Natural enemies of the pear psylla *Cacopsylla pyri* in treated vs untreated pear orchards in Antalya, Turkey. Phytoparasitica, 32, 295–304.

FAOSTAT [WWW Document]. (2020). URL http://www.fao.org/faostat/en/#data/QC [accessed on 2020].

Ge, Y., Liu, P., Zhang, L., Snyder, W.E., Smith, O.M. & Shi, W. (2020) A sticky situation: honeydew of the pear psylla disrupts feeding by its predator *Orius sauteri*. Pest Management Science, 76, 75–84.

Guerrieri, E. & Noyes, J.S. (2009) A review of the European species of the genus *Trechnites* Thomson (Hymenoptera: Chalcidoidea: Encyrtidae), parasitoids of plant lice (Hemiptera: Psylloidea) with description of a new species. Systematic Entomology, 34, 252–259.

Hamilton, W.D. (1967) Extraordinary sex ratios. A sex-ratio theory for sex linkage and inbreeding has new implications in cytogenetics and entomology. Science (New York, N.Y.), 156, 477–488.

Heimpel, G.E. & Collier, T.R. (1996) The evolution of host-feeding behaviour in insect parasitoids. Biol. Rev., 71, 373–400.

Herard, F. (1985) Analysis of parasite and predator populations observed in pear orchards infested by Psylla pyri (L.) (Hom.LJ: Psyllidae) in France. Agronomie, 5, 773–778.

Jaworska, K., Olszak, R.W. & Zajac, R.Z. (1996) Parasitization rate on the larvae of pear (*Cacopsylla pyri*) in orchards with differing intensity of chemical control. Acta Horticulturae, 422, 334–335.

Jerinic-Prodanovic, D., Protic, L. & Mihajlovic, L. (2010) Predators and parasitoids of *Cacopsylla pyri* (L.) (Hemiptera: Psyllidae) in Serbia. Pesticidi i fitomedicina, 25, 29–42.

Jervis, M.A. & Kidd, N.A.C. (1986) Host-feeding strategies in Hymenopteran parasitoids. Biol. Rev., 61, 395–434.

Jia, Y.-J. & Liu, T.-X. (2018) Dynamic host-feeding and oviposition behavior of an aphid parasitoid *Aphelinus asychis*. BioControl, 63, 533–542.

Lacey, L., Arthurs, S., Horton, D. & Miliczky, G. (2005) Spinosad and granulovirus effects on codling moth. USDA-ARS, Yakima Agricultural Research Laboratory, Wapato, WA, Wapato.

Le Goff, G.J., Lebbe, O., Lohaus, G., Richels, A., Jacquet, N., Byttebier, V., et al. (2019) What are the nutritional needs of the pear psylla *Cacopsylla pyri*? Arthropod-Plant Interactions, 13, 431–439.

Mansfield, S. & Mills, N.J. (2002) Host Egg Characteristics, Physiological Host Range, and Parasitism Following Inundative Releases of *Trichogramma platneri* (Hymenoptera: Trichogrammatidae) in Walnut Orchards. Environmental Entomology, 31, 723–731.

McMullen, R.D. (1966) New Records of Chalcidoid Parasites and Hyperparasites of *Psylla pyricola* Förster in British Columbia. The Canadian Entomologist, 98, 236–239.

Mehrnejad, M.R. & Copland, M.J.W. (2006) Behavioral responses of the parasitoid *Psyllaephagus pistaciae* (Hymenoptera: Encyrtidae) to host plant volatiles and honeydew. Entomological Science, 9, 31–37.

Miliczky, E.R. & Horton, D.R. (2005) Densities of beneficial arthropods within pear and apple orchards affected by distance from adjacent native habitat and association of natural enemies with extra-orchard host plants. Biological Control, 33, 249–259.

Muratori, F.B., Damiens, D., Hance, T. & Boivin, G. (2008) Bad housekeeping: why do aphids leave their exuviae inside the colony? BMC Evolutionary Biology, 8.

Oudeh, B., Kassis, W. & Abu-Tara, R. (2013) Seasonal Activity of the Predator, *Anthocoris nemoralis* (F.) and the Parasitoid, *Trechnites psyllae* (R.) against the Pear Psylla *Cacopsylla pyricola* (F.) (Hemiptera: Psyllidae). Egyptian Journal of Biological Control, 23, 17–23.

Pan, M.-Z., Wang, L., Zhang, C.-Y., Zhang, L.-X. & Liu, T.-X. (2017) The influence of feeding and host deprivation on egg load and reproduction of an aphid parasitoid, *Aphidius gifuensis* (Hymenoptera: Braconidae). Applied Entomology and Zoology, 52, 255–263.

Retan, A. & Peterson, V. (1982) Pear Psylla. Insect answers.

Sanchez, J.A. & Ortin-Angulo, M.C. (2012) Abundance and population dynamics of *Cacopsylla pyri* (Hemiptera: Psyllidae) and its potential natural enemies in pear orchards in southern Spain. Crop Protection, 32, 24–29.

Seemüller, E. & Schneider, B. (2004) ‘Candidatus *Phytoplasma mali*’, ‘Candidatus *Phytoplasma pyri*’ and ‘Candidatus *Phytoplasma prunorum*’, the causal agents of apple proliferation, pear decline and European stone fruit yellows, respectively. International Journal of Systematic and Evolutionary Microbiology, 54, 1217–1226.

Sigsgaard, L., Esbjerg, P. & Philipsen, H. (2006a) Controlling pear psyllids by mass releasing *Anthocoris nemoralis* and *A. nemorum* (Heteroptera: Anthocoridae). Journal of Fruit and Ornamental Plant Research, 14, 10.

Sigsgaard, L., Esbjerg, P. & Philipsen, H. (2006b) Experimental releases of *Anthocoris nemoralis* F. and *Anthocoris nemorum* (L.) (Heteroptera: Anthocoridae) against the pear psyllid *Cacopsylla pyri* L. (Homoptera: Psyllidae) in pear. Biological Control, 39, 87–95.

Tougeron, K., Ilitis, C., Renoz, F., Albitar, L., Hance, T., Demeter, S., Le Goff, G.J. Ecology and biology of the parasitoid *Trechnites insidiosus* and its potential for biological control of pear psyllids. Pest management science.

Unruh, T.R., Westigard, P.H. & Hagen, K.S. (1994) Pear Psylla. In Biological control in the Western United State. DANR Publications, pp. 95–100.

Van Alphen, J.J.M. & Visser, M.E. (1990) Superparasitism as an Adaptive Strategy for Insect Parasitoids. Annual Review of Entomology, 35, 59–79.

Van Den Assem, J., Gijswijt, M.J. & Nübel, B.K. (1982) Characteristics of courtship and mating behaviour used as classificatory criteria in Eulophidae-Tetrastichinae (Hymenoptera), with special reference to the genus *Tetrastichus*. Tijdscrift voor entomologie, 125, 205–219.

Villagra, C.A., Ramirez, C.C. & Niemeyer, H.M. (2002) Antipredator responses of aphids to parasitoids change as a function of aphid physiological state. Animal behaviour, 64, 677–683.

Walker, A.M. & Hoy, M.A. (2003) Responses of *Lipolexis oregmae* (Hymenoptera: Aphidiidae) to Different Instars of Toxoptera citricida (Homoptera: Aphididae). Journal of Economic Entomology, 96, 1685–1692.

Zhukovskaya, M., Yanagawa, A. & Forschler, B.T. (2013) Grooming Behavior as a Mechanism of Insect Disease Defense. Insects, 4, 609–630.

