## Supplementary material for "Impact of the pear psyllid *Cacopsylla pyri* host instar on the behavior and fitness of the parasitoid *Trechnites insidious*": supplementary data 1.docx

*Z-values of the Tukey’s multiple comparison test between each developmental instar for the number of antennal contacts*

|  | **Z-value** | **Pr(>\|z\|)** |
| --- | --- | --- |
| **Instar 2 - Instar 1** | -4.73 | <0.001*** |
| **Instar 3 - Instar 1** | 6.46 | <0.001*** |
| **Instar 4 - Instar 1** | 6.31 | <0.001*** |
| **Instar 5 - Instar 1** | -0.82 | 0.92 |
| **Instar 3 - Instar 2** | 10.38 | <0.001*** |
| **Instar 4 - Instar 2** | 10.25 | <0.001*** |
| **Instar 5 - Instar 2** | 3.96 | <0.001*** |
| **Instar 4 - Instar 3** | -0.16 | 1.00 |
| **Instar 5 - Instar 3** | -7.2 | <0.001*** |
| **Instar 5 - Instar 4** | -7.05 | <0.001*** |

*Z-values of the Tukey’s multiple comparison test between each developmental instar for the number of ovipositor insertions*

|  | **Z-value** | **Pr(>\|z\|)** |
| --- | --- | --- |
| **Instar 2 - Instar 1** | -4.27 | <0.001*** |
| **Instar 3 - Instar 1** | 3.03 | 0.02* |
| **Instar 4 - Instar 1** | 4.61 | <0.001*** |
| **Instar 5 - Instar 1** | -4.9 | <0.001*** |
| **Instar 3 - Instar 2** | 6.85 | <0.001*** |
| **Instar 4 - Instar 2** | 8.12 | <0.001*** |
| **Instar 5 - Instar 2** | -0.75 | 0.94 |
| **Instar 4 - Instar 3** | 1.66 | 0.45 |
| **Instar 5 - Instar 3** | -7.34 | <0.001*** |
| **Instar 5 - Instar 4** | -8.53 | <0.001*** |

*Z-values of the Tukey’s multiple comparison test between each developmental instar for the host acceptance*

|  | **Z-value** | **Pr(>\|z\|)** |
| --- | --- | --- |
| **Instar 2 - Instar 1** | -0.15 | 1.00 |
| **Instar 3 - Instar 1** | -0.53 | 0.98 |
| **Instar 4 - Instar 1** | -0.07 | 1.00 |
| **Instar 5 - Instar 1** | -3.35 | 0.01** |
| **Instar 3 - Instar 2** | -0.33 | 1.00 |
| **Instar 4 - Instar 2** | 0.1 | 1.00 |
| **Instar 5 - Instar 2** | -2.99 | 0.02* |
| **Instar 4 - Instar 3** | 0.53 | 0.98 |
| **Instar 5 - Instar 3** | -3.16 | 0.01* |
| **Instar 5 - Instar 4** | -3.79 | 0.001** |

*Z-values of the Tukey’s multiple comparison test between each developmental instar for the time walking*

|  | **Z-value** | **Pr(>\|z\|)** |
| --- | --- | --- |
| **Instar 2 - Instar 1** | -1.38 | 0.64 |
| **Instar 3 - Instar 1** | 1.1 | 0.81 |
| **Instar 4 - Instar 1** | 0.71 | 0.95 |
| **Instar 5 - Instar 1** | 1.98 | 0.28 |
| **Instar 3 - Instar 2** | 2.48 | 0.10. |
| **Instar 4 - Instar 2** | 2.09 | 0.22 |
| **Instar 5 - Instar 2** | 3.36 | 0.007** |
| **Instar 4 - Instar 3** | -0.39 | 0.99 |
| **Instar 5 - Instar 3** | 0.88 | 0.91 |
| **Instar 5 - Instar 4** | 1.26 | 0.71 |

*Z-values of the Tukey’s multiple comparison test between each developmental instar for the time resting*

|  | **Z-value** | **Pr(>\|z\|)** |
| --- | --- | --- |
| **Instar 2 - Instar 1** | 1.88 | 0.33 |
| **Instar 3 - Instar 1** | -1.58 | 0.51 |
| **Instar 4 - Instar 1** | -1.94 | 0.30 |
| **Instar 5 - Instar 1** | -1.95 | 0.29 |
| **Instar 3 - Instar 2** | -3.46 | 0.005** |
| **Instar 4 - Instar 2** | -3.81 | 0.001** |
| **Instar 5 - Instar 2** | -3.83 | 0.001** |
| **Instar 4 - Instar 3** | -0.35 | 1.00 |
| **Instar 5 - Instar 3** | -0.37 | 1.00 |
| **Instar 5 - Instar 4** | -0.02 | 1.00 |

*Z-values of the Tukey’s multiple comparison test between each developmental instar for the number of mummies*

|  | **Z-value** | **Pr(>\|z\|)** |
| --- | --- | --- |
| **Instar 2 - Instar 1** | 0.71 | 0.94 |
| **Instar 3 - Instar 1** | 5.61 | <0.001*** |
| **Instar 4 - Instar 1** | 4.9 | <0.001*** |
| **Instar 5 - Instar 1** | -0.02 | 1.00 |
| **Instar 3 - Instar 2** | 5.26 | <0.001*** |
| **Instar 4 - Instar 2** | 4.48 | <0.001*** |
| **Instar 5 - Instar 2** | -0.02 | 1.00 |
| **Instar 4 - Instar 3** | -1.04 | 0.8 |
| **Instar 5 - Instar 3** | -0.02 | 1.00 |
| **Instar 5 - Instar 4** | -0.02 | 1.00 |

*Z-values of the Tukey’s multiple comparison test between each developmental instar for the host suitability*

|  | **Z-value** | **Pr(>\|z\|)** |
| --- | --- | --- |
| **Instar 2 - Instar 1** | 2.43 | 0.1 |
| **Instar 3 - Instar 1** | 3.42 | 0.005** |
| **Instar 4 - Instar 1** | 1.89 | 0.32 |
| **Instar 5 - Instar 1** | -1.23 | 0.73 |
| **Instar 3 - Instar 2** | 0.6 | 0.97 |
| **Instar 4 - Instar 2** | -0.9 | 0.90 |
| **Instar 5 - Instar 2** | -3.8 | 0.001** |
| **Instar 4 - Instar 3** | -1.8 | 0.37 |
| **Instar 5 - Instar 3** | -5.2 | <0.001*** |
| **Instar 5 - Instar 4** | -3.53 | 0.004** |

*Z-values of the Tukey’s multiple comparison test between each developmental instar for the tibia size*

|  | **Z-value** | **Pr(>\|z\|)** |
| --- | --- | --- |
| **Instar 4 - Instar 2** | 3.18 | 0.007** |
| **Instar 3 - Instar 2** | 3.1 | 0.009** |
| **Instar 1 - Instar 2** | 2.40 | 0.07. |
| **Instar 3 - Instar 4** | 0.06 | 0.99993 |
| **Instar 1 - Instar 4** | -0.07 | 0.99987 |
| **Instar 1 - Instar 3** | -0.1 | 0.99959 |

*Z-values of the Tukey’s multiple comparison test between each developmental instar for the developmental time*

|  | | | **Z-value** | **Pr(>\|z\|)** |
| --- | --- | --- | --- | --- |
| **Instar 4 - Instar 2** |  |  | -2.71 | 0.03* |
| **Instar 3 - Instar 2** |  |  | -2.10 | 0.15 |
| **Instar 1 - Instar 2** |  |  | 4.14 | <0.001*** |
| **Instar 3 - Instar 4** |  |  | 0.90 | 0.80 |
| **Instar 1 - Instar 4** |  |  | 8.07 | <0.001*** |
| **Instar 1 - Instar 3** |  |  | 7.32 | <0.001*** |
